# Imaging brain vascular function in Cystic Fibrosis: an MRI study of cerebral blood flow and brain oxygenation

**DOI:** 10.1101/2024.02.25.581905

**Authors:** HL Chandler, M Germuska, TM Lancaster, C Xanthe, C O’leary, S Stirk, K Murphy, C Metzler-Baddeley, RG Wise, J Duckers

## Abstract

Cystic fibrosis (CF) is a progressive inherited disorder that primarily affects the lungs. With recent breakthroughs in effective treatments for CF that increase life-expectancy, a higher prevalence of age-related comorbidities have been reported including cardiovascular disease, stroke and cognitive decline. Despite the known relationship between cardiovascular health and cerebrovascular function, very little is known about brain blood flow and oxygen metabolism in patients with CF (PwCF). In 14 PwCF and 56 healthy age / sex matched controls, we used pseudo-continuous arterial spin labelling (pCASL) to quantify cerebral perfusion in grey-matter and T_2_-Relaxation-Under-Spin-Tagging (TRUST) to estimate global oxygen extraction fraction (OEF) and cerebral metabolic rate of oxygen consumption (CMRO_2_). Compared to healthy controls, PwCF showed elevated CMRO_2_ (*p =* 0.015). There were no significant between-group differences in grey-matter CBF (*p =* 0.342), or whole brain OEF (*p =* 0.091). However, regional analysis showed certain areas with higher CBF in PwCF (*p* < .05, FDR). This is the first study to characterise cerebrovascular function and brain oxygen metabolism in PwCF. Our findings highlight the need for early cardiovascular monitoring procedures to help maintain cerebrovascular function and combat accelerated aging effects in the brains of PwCF.

## Introduction

Cystic fibrosis (CF) is a progressive inherited multi-system disease arising from a mutation in the transmembrane conductance regulator (CFTR) gene (1). Dysfunction in the CFTR protein causes a thick mucus to build up in the lungs, leading to a series of pathological effects on the respiratory and circulatory system, including recurrent infection, hypoxia, widespread inflammation, and vascular dysfunction (2-5). These effects can have a significant long-term impact on the brain, especially given the crucial role of the cerebrovascular system in satisfying the brain’s need for uninterrupted delivery of oxygen and nutrients.

Patients with CF (PwCF) frequently present with brain related symptoms that affect mood, memory, attention, and executive function (6, 7), however, the underlying cause of these effects remains unknown. Cardiovascular risk factors are disproportionally prevalent earlier in the lifespan in PwCF, including CF related diabetes, arterial stiffness, and hypertension (8). There is increasing interest in the heart-brain relationship in neurodegenerative disease research, specifically how cardiovascular risk leads to altered brain function and cognitive decline. Arterial stiffness is prevalent in children with CF (9), where early cardiovascular risk factors may lead to disrupted brain vascular function, and eventually downstream cognitive decline. However, to date there is a stark lack of research into brain health and function of patients with CF.

One magnetic resonance imaging (MRI) study of 5 PwCF and 15 healthy controls revealed differences in grey-matter density and T_2_ relaxation times (6). The authors suggested that these brain differences may reflect hypoxia and neuronal damage (10). Hypoxia is thought to occur when the body’s organs / tissues are starved of oxygen. Hypoxia in the brain can have a significant impact on cognition, including memory, learning and attention (11), and may lead to premature aging and neurodegeneration (12). However, it is unclear how subtle effects of hypoxia (and the type of hypoxia) on the brain in CF would impact cognition, and whether this would be reflected as altered brain vascular function.

In other respiratory diseases, such as chronic obstructive pulmonary disease (COPD), patients present with reduced cerebral blood flow (13) and vascular pathologies including strokes and microbleeds (14). However, COPD is typically diagnosed later in life when environmental factors also contribute to an increased prevalence of cardiovascular comorbidities. Therefore, reduced CBF may not be expected in PwCF. Evidence suggests that children with cystic fibrosis show elevations in the partial pressure of carbon dioxide (PaCO_2_) (15). Elevated PaCO_2_ has vasodilatory effects, which could result in increased CBF (16, 17). A fall in PaO_2_, if below 40-45mmHg, could also result in increased CBF (17).

Hypoxia causes a disruption in cerebral autoregulation, angiogenesis, an increase in the production of endothelial derived nitric oxide (NO), all of which are also evidenced to increase CBF (17). Given the known coupling between CBF and brain O_2_ consumption, and the link between hypoxia and vascular function in other respiratory diseases (18), it is plausible to suggest that PwCF would show altered cerebrovascular function.

To the best of our knowledge, no research has yet characterised cerebral blood flow or cerebral metabolic oxygen consumption (CMRO_2_) in PwCF. Assessment of vascular function in CF is vital, particularly at the microvascular level where effective delivery, exchange, and metabolism of oxygen is key to maintaining mitochondrial oxygen homeostasis and thus healthy neuronal function (19, 20). In the present study we hypothesise that PwCF will have altered cerebrovascular function compared to healthy age / sex matched controls, specifically in relation to CBF and CMRO_2_. We used pseudo-continuous arterial spin labelling (pCASL) to quantify grey-matter cerebral blood flow (gmCBF), and T_2_-Relaxation-Under-Spin-Tagging (TRUST) MRI to estimate global oxygen extraction fraction (OEF) and the cerebral metabolic rate of oxygen consumption (CMRO_2_) (21). By characterising these cerebrovascular parameters, we will have a better understanding of the effects that CF has on brain tissue function and thus provide a first step towards developing strategies that monitor cardiovascular disease, protect vascular function, and prevent downstream cognitive decline.

## Methods

### Participants

Fourteen patients with a diagnosis of CF and 56 age / sex matched healthy controls took part in the present brain imaging study (see *Table 1*.). Eligibility criteria of the patients included a primary diagnosis of CF, primary school level education and above, and ability to provide informed consent. Exclusion criteria included any MRI contraindications, existing neurological or neuropsychiatric disease (including dementia), any vascular complications of diabetes, requiring supplemental oxygen (long term or overnight), smoking, pregnancy, current infection, or hepatic failure.

**Table 1.** Clinical and demographic data for the study population. Results include means and standard deviations. We also report patient group characteristics, including a series of blood assays. ^a^ Sex differences tested via chi-squared test; ^b^ age and BMI differences are calculated via independent samples t-test. Neuropsychology / cognitive data in the table is presented with group means, and range in parentheses.

|  | Patients | Controls | *p*-value |
| --- | --- | --- | --- |
| Sex (M / F) | 11 / 3 | 42 / 14 | 1<sup>a</sup> |
| Age (years) | $34.57 \pm 10.92$ | $34.48 \pm 11.78$ | 0.978<sup>b</sup> |
| BMI (Kg/m<sup>2</sup>) | $24.83 \pm 3.78$ | $25.21 \pm 3.33$ | 0.712<sup>b</sup> |
| Blood: HbA1C (mmol/L) | $52.08 \pm 28.61$ | | |
| Blood: Hb (g/L) | $149.07 \pm 13.77$ | | |
| Blood: Ferritin (ng/mL) | $42.92 \pm 32.04$ | | |
| Blood: HDL (mmol/L) | $1.38 \pm 0.30$ | | |
| Blood: LDL (mmol/L) | $2.64 \pm 0.42$ | | |
| Telomere TL (kb) | $5.12 \pm 1.32$ | | |
| Test of Premorbid Functioning - Full Scale IQ | 99.73 (82-114) |  |  |
| Wechsler Memory Scale: Logical Memory, Immediate Recall | 9.81 (5-15) |  |  |
| Wechsler Memory Scale: Logical Memory, Delayed Recall | 9.00 (4-13) |  |  |
| Ray Complex Figure: Immediate Score | 54.27 (34-80) |  |  |
| Ray Complex Figure: Delayed Score | 51.81 (31-68) |  |  |
| Wechsler Memory Scale: Working Memory Index | 102.09 (89-128) |  |  |
| Wechsler Memory Scale: Performance Scale Index | 108.73 (92-124) |  |  |

Written informed consent was obtained from all patients and healthy volunteers who took part in the two studies. The patient arm of the study was approved by the NHS Research Ethics Committee (NHS HRA REC reference 20/WA/0068), Wales, UK and both patient / healthy volunteer studies obtained ethical approval from Cardiff University’s Ethics Committee (add both ethics committee numbers), School of Psychology, Cardiff, UK, in accordance with the Declaration of Helsinki.

## Data collection

### Clinical data and behavioural assessments

PwCF underwent neuropsychological testing and venous blood samples were collected and tested for C-Reactive Protein (CRP), ferritin, hemoglobin A1C (HBA1C), hemoglobin (Hb), fasting lipids (high-density lipoprotein (HDL); low-density lipoprotein (LDL). Telomere data collected from blood were also analysed for the PwCF. Patient blood samples were collected within 3 weeks of their MRI scan at CUBRIC.

### MRI acquisition

All MRI data were collected on a Siemens Prisma 3T MRI scanner (*Siemens Healthineers, Erlangen, Germany)*, with a 32-channel receive only head coil.

### Structural scans

A magnetisation-prepared rapid acquisition with gradient echo (MPRAGE) T1-weighted scan was used for registration, brain segmentation purposes, and to compare gross structural differences between groups including intracranial volume, grey-matter volume, white-matter volume (matrix 165 x 203 x 197, 1 mm isotropic resolution, TR/TE = 2100/3.24ms).

### Functional MRI scans

To quantify gmCBF, an in-house single post label delay (PLD) PCASL sequence was collected (110 volumes; TR/TE= 4600ms / 11ms; slices = 22; Slice thickness = 5mm; PLD = 2000ms; tag duration = 1800ms; GRAPPA acceleration factor = 2). An M0 was collected for calibration and thus CBF quantification (TR/TE= 6000ms / 11ms; slices = 22; Slice thickness = 5mm; GRAPPA acceleration factor = 2).

A T2-Relaxation-Under-Spin-Tagging (TRUST) MRI sequence was acquired to estimate global venous oxygenation and OEF (TR / TE = 3000ms / 3.9ms, slice thickness = 5mm, eTE = 0ms, 40ms, 80ms, 160ms, GRAPPA acceleration factor = 3) from venous blood in the sagittal sinus. We further collected a T1 inversion recovery sequence to quantify the T_1_ of venous blood to calculate [Hb] (22), necessary for estimating OEF and CMRO_2_ (ΔTR / TE = 150ms / 22ms, flip angle = 90 degrees, and GRAPPA acceleration factor = 2; with 960 acquisitions > 16 repeats of 60 measurements).

## Data / statistical analysis

### Structural MRI

The T1-weighted structural MRI was segmented and reconstructed via Freesurfer version 7.1.1 (https://surfer.nmr.mgh.harvard.edu/fswiki/ReleaseNotes). Briefly, the FreeSurfer recon-all pipeline processes structural MRI data, then removes non-brain tissues, and segments the brain into grey matter, white matter, and cerebrospinal fluid. Subsequently, it reconstructs the cortical surface, defining gyri and sulci. Cortical parcellation divides the surface into distinct regions, facilitating region-specific investigations. We consider intracranial volume (ICV; mm3), grey-matter volume (ml), and white-matter volume (ml) based on the method outlined by Fischl et al. (23). All T1-weighted images were further delineated and labelled using their cytoarchitectural boundaries, according to the Desikan-Killiany Atlas (24), which was used for regional analysis of the gmCBF (see Methods: Regional CBF).

### Quantitative Functional MRI

#### GM CBF and Global tissue oxygenation

We used the FSL Bayesian Inference for Arterial Spin Labeling MRI *(BASIL)* toolbox (FSL 6.0.1) to analyse our CBF data implementing motion correction, partial volume estimation (PVE), and CBF quantification using the M0 with CSF as a reference for calibration to get flow into absolute units (25, 26). We created binarized grey-matter masks with a PV threshold of 0.8 (80%) to create a new gmCBF PV corrected map. The average gmCBF was extracted from this map and used in the CMRO_2_ calculation. The T_2_ of blood was derived by non-linear least squared fitting of a mono-exponential equation to the TRUST difference data. This was as a function of the effective echo times and T_2_. TRUST analysis involved using the two most intense voxels from a sagittal sinus ROI and venous oxygenation (Yv) was estimated by inverting the relationship between Yv, [Hb] and the T_2_ (21). OEF was calculated as OEF = ((SaO_2_-SvO_2_)/SaO_2_), and from here we estimated global CMRO_2_ = gmCBF * CaO_2_ = ([Hb] * 1.34 * SaO_2_) * OEF * 39.34. We used an assumed SaO_2_ value of 98% for the healthy cohort and 98% for the patient cohort (supported by SpO_2_ data taken from each patient during their research visit at clinic). Mean regional gmCBF values were further assayed across cyto-architecturally defined regions characterised within the Desikan Killany atlas (24).

## Statistical analysis

We performed a series of simultaneous multiple linear regressions to assess the relationship between 1) a series of outcomes (γ) including 1a) assays of whole brain cortical macrostructure (including intracranial volume, grey and white matter volume (ml), and 1b) whole brain (CBF, CMRO_2_, OEF) & regional (CBF only measure assessed regionally) and 2) independent variables including the presence of a CF diagnosis (χ_1_) and covariates of no interest including age (χ_2_) and sex (χ_3_), which provide explanatory variance in macrostructure and cerebrovascular estimates (27). Outliers for dependent variables were identified and iteratively removed using the outlier labelling rule, an established recommendation for data that may exert disproportional impact on regression coefficients (28). Briefly, each dependent variable was centered around the median and all points outside 1.5 × IQR (Interquartile Range) (Q3 – Q1) were labelled as outliers and removed from consideration. There were no significant changes to any results before / after the removal of outliers for any dependent variable considered. For regional CBF mapping, the p-values corresponding to CF coefficients for 68 cyto-architecturally distinct regions (34, per hemisphere) were corrected via the false discovery rate (FDR).

## Results

Patient characteristics are presented in *Table 1*. Analysis included fourteen patients (mean age ± SD = 34.57 ± 10.92, 11/3 male/female) and 55 healthy controls (mean age ± SD = 34.48 ± 11.78, 42/14 male/female).

## Imaging analysis

### Structural

Results show no statistically significant between-group differences in ICV, grey-matter volume or white-matter volume (see Table 2).

**Table 2.** Independent sample t-tests between PwCF and healthy controls looking at brain structure. Abbreviations: standard deviation (SD); intracranial volume (ICV).

| Global Metric | Patients | Controls | *t*-statistic | *p*-value |
| --- | --- | --- | --- | --- |
| ICV (ml) | 1586.31<br>± 128.02 | 1617.90<br>± 168.13 | -0.829 | 0.410 |
| Grey-matter<br>volume (ml) | 684.02 ±<br>59.99 | 693.22 ±<br>62.91 | 0.508 | 0.612 |
| White-matter<br>volume (ml) | 491.04 ±<br>46.43 | 483.10 ±<br>54.74 | 0.551 | 0.587 |

### Global CBF, OEF and CMRO_2_

There were significant increases in CMRO_2_ (PwCF vs. controls: 190.46 vs.

169.36μmol/100 g/min, *β* = 21.43 ± 8.53, *p* = 0.015) in PwCF compared to healthy controls. There were no significant between-group differences in grey-matter CBF (PwCF vs. controls: 65.32 vs. 62.93ml/100 g/min, *β* = 2.68 ± 2.8, *p* = .342) or OEF (PwCF vs. controls: 0.41 vs. 0.37, *β* = 0.03 ± 0.02, *p* = 0.091) (*Figure 1*).

**Figure 1.**
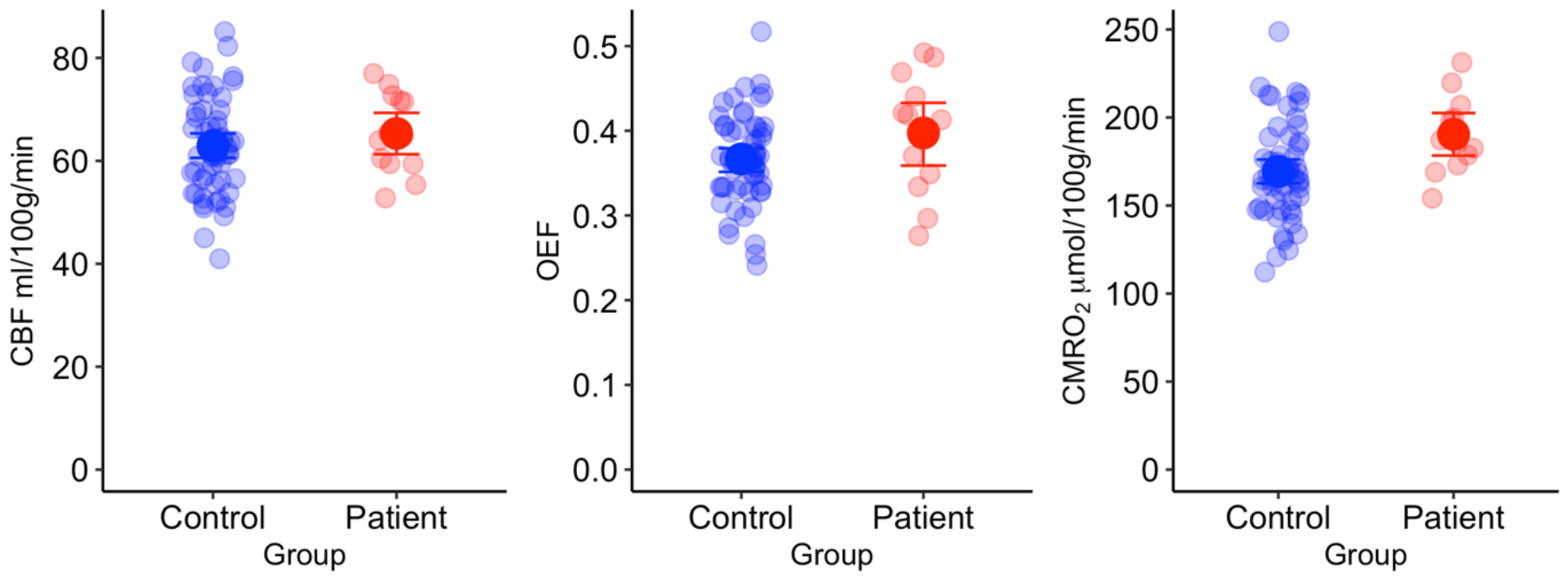
Results reveal significant increases in global CMRO_2_ (*p* =.015) in the patient group compared to age and sex matched controls. Our results showed no significant differences in whole grey-matter CBF (*p* =.342) or OEF (*p* =.091). Error bars reflect 95% confidence intervals. Abbreviations: cerebral blood flow (CBF); oxygen extraction fraction (OEF); cerebral metabolic rate of oxygen consumption (CMRO_2_).

**Figure 2.**
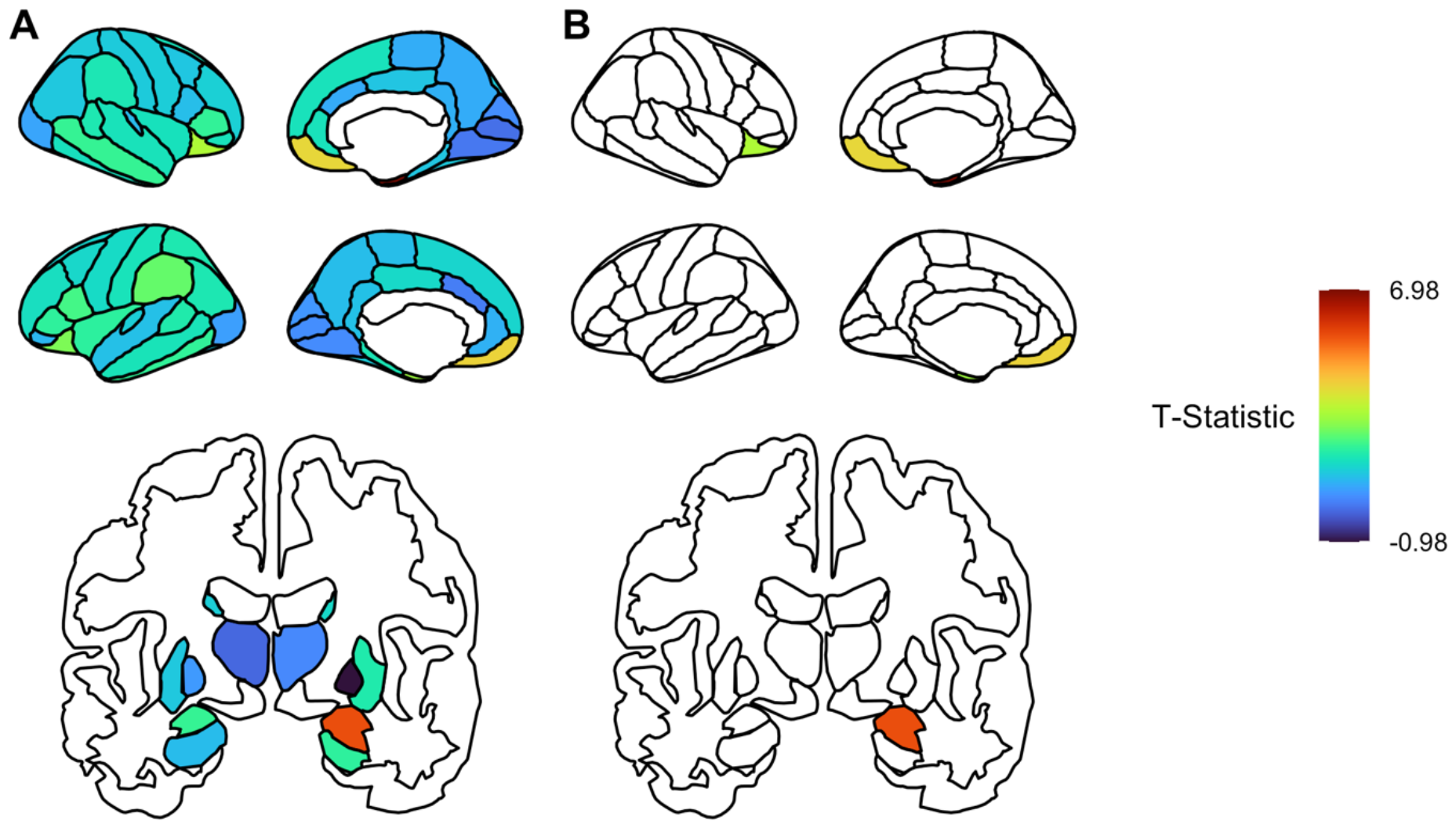
Group level contrast (PwCF > Healthy Controls) for differences in regional grey-matter CBF plotted as the t-statistic both uncorrected (a) and FDR corrected (b). The group level differences are plotted across the cortex (upper rows, sagittal) and subcortical regions (lower rows, coronal) separately.

### Regional CBF analysis

While we observed no significant difference in global gm-CBF between PwCF and controls, our regression analysis identified several cortical regions where PwCF had significantly higher gm-CBF compared to the healthy controls, following correction for false discovery rate (FDR, Benjamini and Hochberg), see *Table 3*. There were no regions where PwCF showed significantly lower CBF compared to controls.

**Table 3.** Beta estimates (and standard error of estimate) for association between PwCF diagnosis and regional gm-CBF features. Abbreviations: standard error (SE); false discovery rate (FDR).

| Region of Interest | Hemisphere | Beta Estimate<br>(ml/100g/min) | SE | *t*-statistic | *P* | *P* FDR |
| --- | --- | --- | --- | --- | --- | --- |
| Entorhinal | L | 5.070 | 1.726 | 2.938 | 0.005 | 0.041 |
| Entorhinal | R | 10.05 | 1.44 | 6.99 | <.001 | <.001 |
| Amygdala | R | 10.76 | 1.94 | 5.54 | <.001 | <.001 |
| Temporal pole | L | 6.62 | 2.168 | 3.052 | 0.003 | 0.037 |
| Temporal pole | R | 10.14 | 1.95 | 5.19 | <.001 | <.001 |
| Medial<br>orbitofrontal | L | 7.94 | 2.00 | 3.96 | <.001 | 0.004 |
| Medial<br>orbitofrontal | R | 10.56 | 2.72 | 3.89 | <.001 | 0.004 |
| Lateral<br>orbitofrontal | R | 6.517 | 2.076 | 3.138 | 0.003 | 0.034 |
| Nucleus<br>accumbens | R | 7.659 | 2.538 | 3.017 | 0.004 | 0.037 |

## Discussion

This study is the first to evidence cerebrovascular and brain oxygen metabolism differences in PwCF using MRI. Our results show that PwCF have increased regional perfusion and cerebral oxygen metabolism, in the absence of whole grey-matter cerebral perfusion differences, oxygen extraction fraction, or macrostructural differences. Furthermore, the absence of significant grey matter volume differences between PwCF and controls suggests that these changes in tissue physiology may precede atrophy in patients and might be a target for accelerated aging strategies.

Despite the evidenced association between increased CBF and CMRO_2_ in a state of hypoxia (29), our patients SpO_2_ values were within the expected range for a healthy population (98%), and no patients were on oxygen therapy, suggesting that our group were not displaying signs of hypoxia at the time of the scan. However, hypoxia is known to have a negative impact on cognitive function including attention, learning and memory (11, 30-32), and may eventually lead to neurodegenerative disease (12). Several mechanisms have been suggested to explain the link between hypoxia and cognitive decline, including reduced ATP through glycolysis, oxidative stress, calcium overload, and mitochondrial injury (11, 33).

Another explanation for observing an increase in CBF and CMRO_2_ could be group differences in PaCO_2_ / PaO_2_. As we were unable to collect direct data on the partial pressure of CO_2_ or O_2_, we are unable to ascertain whether these factors explain our results. We suggest that future research studies would benefit from collecting this data where possible, especially studies with a focus on brain vascular function in pulmonary disease.

There have been significant breakthroughs in treatments for CF, including CFTR modulators, that have significantly extended patients’ life expectancy. However, this appears to have come at a cost of a higher prevalence of age-related cardiovascular comorbidities.

There is increasing interest in understanding how to manage the aging CF population, especially with the increase in risk factors such as inflammation, CF related diabetes and obesity, all of which can cause significant vascular pathologies. Preserving cardiovascular health may help to prevent accelerated aging effects in the brain including inflammation, mitochondrial dysfunction, stroke, and early cognitive decline. Cardiovascular comorbidities are highly prevalent in PwCF (34) with manifestations including pulmonary hypertension, atherosclerosis, cardiomyopathies, and endothelial cell dysfunction (5, 35, 36). These effects are also well known to have several damaging effects on the brain, including dysfunction in cerebral blood flow, tissue oxygenation, and increased risk of stroke, and cognitive decline (37-41). However, evidence from patients with COPD with cardiovascular risk suggests that cerebral perfusion is reduced (13), rather than increased.

Our results may also be explained by altered endothelial cell function in PwCF. Emerging evidence suggests that CFTR is expressed in endothelial cells and linked to endothelial cell dysfunction (36). Endothelial cells have many functions, including supporting blood vessel formation, maintaining the blood-brain barrier, and regulation of vascular tone (42). This is of particular importance in PwCF who show a higher prevalence of endothelial cell dysfunction compared to the general population, as well as heightened levels of vascular inflammation, oxidative stress, and vascular stiffness (43-45). Future research could benefit from exploring regional alteration in OEF and CMRO_2_ via dual-calibrated MRI protocols (and alike). With these approaches, we can also estimate the partial pressure of oxygen at the mitochondrial level (46), which may be useful in assessing more localised microvascular alterations in PwCF. Our results could also reflect a dysfunction in the electron transport chain, specifically at complex IV and V, where a dysfunction / inefficient use of oxygen utilisation could be reflected as an increase in CMRO_2_ to maintain ATP production.

Our structural analysis showed no statistically significant differences in grey matter volume, white matter volume or intercranial volume. It has been suggested that PwCF show premature / accelerated aging, which may be linked to cognitive changes as patients age. One study has shown structural brain changes in PwCF (6). However, in our study, we identify vascular changes in the absence of structural changes. This suggests that vascular alterations may precede structural neurodegeneration / atrophy, a finding we have observed in other clinical cohorts (46).

Our results should be considered in the light of the following limitations. First, our interpretation is limited to global OEF and CMRO_2_ estimates. With regional measures we would be able to determine more localised vascular differences in PwCF vs healthy controls. Second, our CBF estimates are based on ASL that only provides reliable perfusion measures in grey-matter, not white matter. Third, our study would have been strengthened from collecting overlapping cognitive data in the patients and controls, to assess between-group differences in cognition in relation to vascular measures. Fourth, we are limited by the small patient sample size. With a higher number of patients, we would be powered to look at associations between clinical markers and the vascular IDPs. With strict inclusion / exclusion criteria, the patient cohort were well controlled for cardiovascular and neurological / sever neuropsychiatric disease, all of which could present confounds for our results. Furthermore, our healthy control sample is large and well matched to our patient group for age and sex characteristics.

Our study is the first to characterise cerebrovascular function and brain oxygen metabolism in CF. Given the high prevalence of cardiovascular comorbidities in an aging CF population, and the link between heart health and brain function, there is a stark absence of research in this area. Our work highlights the need for strategic planning for monitoring cardiovascular health in CF, with the aim of preserving brain function and preventing early aging. Our research also provides a basis for developing future therapies that may help to prevent / alleviate downstream cognitive decline.

## Acknowledgements

This study was funded by Welsh Government Pathway to Portfolio funding. HLC & KM: SRF Wellcome [WT224267]. CMB is funded by a National Institute for Health and Care Research (NIHR) and Health and Care Research Wales (HCRW) Advanced Research Fellowship (NIHR-FS(A)-2022). The healthy controls from this study were recruited for a larger study in CUBRIC funded by a Wellcome Strategic Award [104943/Z/14/Z].

## Data Availability

The data for the healthy control participants were acquired as part of the Welsh Advanced Neuroimaging Dataset (McNabb et al., 2024). Data is available in BIDS format from https://git.cardiff.ac.uk/cubric/wand.

## References

1. Batten J. Cystic Fibrosis: A Review. Br J Dis Chest. 1965;59:1–9.

2. Petrocheilou A, Moudaki A, Kadis AG. Inflammation and Infection in Cystic Fibrosis: Update for the Clinician. Children (Basel). 2022;9(12).

3. Montgomery ST, Mall MA, Kicic A, Stick SM, Arest CF. Hypoxia and sterile inflamma-on in cystic fibrosis airways: mechanisms and potential therapies. Eur Respir J. 2017;49(1).

4. Patergnani S, Vitto VAM, Pinton P, Rimessi A. Mitochondrial Stress Responses and “Mito-Inflammation” in Cystic Fibrosis. Front Pharmacol. 2020;11:581114.

5. Poore TS, Taylor-Cousar JL, Zemanick ET. Cardiovascular complications in cystic fibrosis: A review of the literature. J Cyst Fibros. 2022;21(1):18–25.

6. Roy B, Woo MS, Vacas S, Eshaghian P, Rao AP, Kumar R. Regional brain tissue changes in patients with cystic fibrosis. J Transl Med. 2021;19(1):419.

7. Chadwick HK, Abbott J, Hurley MA, Dye L, Lawton CL, Mansfield MW, et al. Cystic fibrosis-related diabetes (CFRD) and cognitive function in adults with cystic fibrosis. J Cyst Fibros. 2022;21(3):519–28.

8. Hull JH, Garrod R, Ho TB, Knight RK, Cockcroft JR, Shale DJ, et al. Increased augmentation index in patients with cystic fibrosis. Eur Respir J. 2009;34(6):1322–8.

9. Buehler T, Steinmann M, Singer F, Regamey N, Casaulta C, Schoeni MH, et al. Increased arterial stiffness in children with cystic fibrosis. Eur Respir J. 2012;39(6):1536–7.

10. Yao L, Kan EM, Lu J, Hao A, Dheen ST, Kaur C, et al. Toll-like receptor 4 mediates microglial activation and production of inflammatory mediators in neonatal rat brain following hypoxia: role of TLR4 in hypoxic microglia. J Neuroinflammation. 2013;10:23.

11. Wang X, Cui L, Ji X. Cognitive impairment caused by hypoxia: from clinical evidences to molecular mechanisms. Metab Brain Dis. 2022;37(1):51–66.

12. Daulatzai MA. Death by a thousand cuts in Alzheimer’s disease: hypoxia--the prodrome. Neurotox Res. 2013;24(2):216–43.

13. Wijnant SRA, Bos D, Brusselle G, Grymonprez M, Rietzschel E, Vernooij MW, et al. Comparison of cerebral blood flow in subjects with and without chronic obstructive pulmonary disease from the population-based Rotterdam Study. BMJ Open. 2021;11(12):e053671.

14. Lahousse L, Vernooij MW, Darweesh SK, Akoudad S, Loth DW, Joos GF, et al. Chronic obstructive pulmonary disease and cerebral microbleeds. The Rotterdam Study. Am J Respir Crit Care Med. 2013;188(7):783–8.

15. Kraemer R, Latzin P, Pramana I, Ballinari P, Galla S, Frey U. Long-term gas exchange characteristics as markers of deterioration in patients with cystic fibrosis. Resp Res. 2009;10.

16. Ainslie PN. Have a safe night: intimate protection against cerebral hyperperfusion during REM sleep. J Appl Physiol. 2009;106(4):1031–3.

17. Ainslie PN, Ogoh S. Regulation of cerebral blood flow in mammals during chronic hypoxia: a matter of balance. Exp Physiol. 2010;95(2):251–62.

18. Kent BD, Mitchell PD, McNicholas WT. Hypoxemia in patients with COPD: cause, effects, and disease progression. Int J Chronic Obstr. 2011;6:199–208.

19. Claassen J, Thijssen DHJ, Panerai RB, Faraci FM. Regulation of cerebral blood flow in humans: physiology and clinical implications of autoregulation. Physiol Rev. 2021;101(4):1487–559.

20. Silverman A, Petersen NH. Physiology, Cerebral Autoregulation. StatPearls. Treasure Island (FL)2023.

21. Lu H, Ge Y. Quantitative evaluation of oxygenation in venous vessels using T2-Relaxation-Under-Spin-Tagging MRI. Magn Reson Med. 2008;60(2):357–63.

22. Xu F, Li W, Liu P, Hua J, Strouse JJ, Pekar JJ, et al. Accounting for the role of hematocrit in between-subject variations of MRI-derived baseline cerebral hemodynamic parameters and functional BOLD responses. Hum Brain Mapp. 2018;39(1):344–53.

23. Fischl B. FreeSurfer. NeuroImage. 2012;62(2):774–81.

24. Desikan RS, Segonne F, Fischl B, Quinn BT, Dickerson BC, Blacker D, et al. An automated labeling system for subdividing the human cerebral cortex on MRI scans into gyral based regions of interest. Neuroimage. 2006;31(3):968–80.

25. Chappell MA, Groves AR, MacIntosh BJ, Donahue MJ, Jezzard P, Woolrich MW. Partial volume correction of multiple inversion time arterial spin labeling MRI data. Magn Reson Med. 2011;65(4):1173–83.

26. Chappell MA, Groves AR, Whitcher B, Woolrich MW. Variational Bayesian inference for a nonlinear forward model. Trans Sig Proc. 2009;57(1):223–36.

27. MacDonald ME, Williams RJ, Rajashekar D, Stafford RB, Hanganu A, Sun H, et al. Age-related differences in cerebral blood flow and cortical thickness with an application to age prediction. Neurobiol Aging. 2020;95:131–42.

28. Hoaglin DC, Iglewicz B. Fine-Tuning Some Resistant Rules for Outlier Labeling. Journal of the American Statistical Associa-on. 1987;82(400).

29. Vestergaard MB, Lindberg U, Aachmann-Andersen NJ, Lisbjerg K, Christensen SJ, Law I, et al. Acute hypoxia increases the cerebral metabolic rate - a magnetic resonance imaging study. J Cereb Blood Flow Metab. 2016;36(6):1046–58.

30. Areza-Fegyveres R, Kairalla RA, Carvalho CRR, Nitrini R. Cognition and chronic hypoxia in pulmonary diseases. Dement Neuropsychol. 2010;4(1):14–22.

31. Riba-Llena I, Alvarez-Sabin J, Romero O, Santamarina E, Sampol G, Maisterra O, et al. Nighome hypoxia affects global cognition, memory, and executive func-on in community-dwelling individuals with hypertension. J Clin Sleep Med. 2020;16(2):243–50.

32. McMorris T, Hale BJ, Barwood M, Costello J, Corbett J. Corrigendum to “Effect of acute hypoxia on cognition: A systematic review and meta-regression analysis” Neurosci. Biobehav. Rev. 74 (2017) 225–232. Neurosci Biobehav Rev. 2019;98:333.

33. Burtscher J, Mallet RT, Burtscher M, Millet GP. Hypoxia and brain aging: Neurodegeneration or neuroprotection? Ageing Res Rev. 2021;68:101343.

34. Saunders B, Ranganathan. dentifying and preventing cardiovascular disease in patients with cystic fibrosis. Nature Cardiovascular Research. 2022;1:187–8.

35. Shah PH, Lee JH, Salvi DJ, Rabbani R, Gavini DR, Hamid P. Cardiovascular System Involvement in Cystic Fibrosis. Cureus. 2021;13(7):e16723.

36. Declercq M, Treps L, Carmeliet P, Witters P. The role of endothelial cells in cystic fibrosis. J Cyst Fibros. 2019;18(6):752–61.

37. Marebwa BK, Adams RJ, Magwood GS, Basilakos A, Mueller M, Rorden C, et al. Cardiovascular Risk Factors and Brain Health: Impact on Long-Range Cortical Connections and Cognitive Performance. J Am Heart Assoc. 2018;7(23):e010054.

38. Moroni F, Ammira-E, Rocca MA, Filippi M, Magnoni M, Camici PG. Cardiovascular disease and brain health: Focus on white matter hyperintensities. Int J Cardiol Heart Vasc. 2018;19:63–9.

39. Dadu RT, Fornage M, Virani SS, Nambi V, Hoogeveen RC, Boerwinkle E, et al. Cardiovascular biomarkers and subclinical brain disease in the atherosclerosis risk in communities study. Stroke. 2013;44(7):1803–8.

40. Harrison SL, Ding J, Tang EY, Siervo M, Robinson L, Jagger C, et al. Cardiovascular disease risk models and longitudinal changes in cognition: a systematic review. PLoS One. 2014;9(12):e114431.

41. Nash DT, Fillit H. Cardiovascular disease risk factors and cognitive impairment. Am J Cardiol. 2006;97(8):1262–5.

42. Seals DR, Jablonski KL, Donato AJ. Aging and vascular endothelial function in humans. Clin Sci (Lond). 2011;120(9):357–75.

43. Reverri EJ, Morrissey BM, Cross CE, Steinberg FM. Inflammation, oxidative stress, and cardiovascular disease risk factors in adults with cystic fibrosis. Free Radic Biol Med. 2014;76:261–77.

44. Eising JB, van der Ent CK, Teske AJ, Vanderschuren MM, Uiterwaal C, Meijboom FJ. Young patients with cystic fibrosis demonstrate subtle alterations of the cardiovascular system. J Cyst Fibros. 2018;17(5):643–9.

45. Kartal Ozturk G, Conkar S, Eski A, Gulen F, Keskinoglu A, Demir E. Evaluation of increased arterial stiffness in pediatric patients with cystic fibrosis by augmentation index and pulse wave velocity analysis. Pediatr Pulmonol. 2020;55(5):1147–53.

46. Chandler HL, Stickland RC, Patitucci E, Germuska M, Chiarelli AM, Foster C, et al. Reduced brain oxygen metabolism in patients with multiple sclerosis: Evidence from dual-calibrated functional MRI. J Cereb Blood Flow Metab. 2023;43(1):115–28.

